# Mechanisms of soil carbon preservation illuminated by model mineral-associated organic matter

**DOI:** 10.64898/2025.12.12.694042

**Authors:** Swetha Sridhar, Xiaodong Gao, Tao Sun, Caroline A. Masiello, Caroline M. Ajo-Franklin

## Abstract

A central control on atmospheric CO_2_ is the stabilization of organic carbon in soils. While there are well-described properties associated with stable soil carbon, such as interactions with minerals and conserved chemical features, determining the mechanisms underlying these properties is challenged by soil complexity and heterogeneity. Here we synthesize a naturally complex, tunable model form of mineral-associated organic matter (LabMAOM) to explore mechanisms of stable soil carbon formation and persistence. We show that mineral association and ambient organic matter synergistically stabilize LabMAOM against microbial decomposition. Moreover, different starting compositions of organic matter yield a conserved final LabMAOM chemical composition consistent with natural soils. Our findings open new mechanistic understanding of stable soil carbon and will be of utility to climate modeling and soil health management.

## Main Text

Soil organic matter is the largest reservoir of terrestrial carbon and an essential component of the long-term management and sequestration of atmospheric CO_2_ (*1*). With more stored carbon than the atmosphere and vegetation combined, soils provide multiple ecosystem services, such as nutrient management and water regulation (*2*–*4*), and play pivotal roles in ecosystem productivity and carbon-climate feedback (*5*). While a portion of this carbon is converted to CO_2_ by microbes, a fraction remains preserved for centennia to millennia in high-density organomineral associates, termed mineral-associated organic matter (MAOM) (*6*–*10*). Due to its long residence time in soils, MAOM is a promising reservoir for soil carbon sequestration that merits further investigation for long-term soil management and restoration (*10*–*12*).

Despite its importance for global soil carbon sequestration, studies on the chemical nature and biological behavior of MAOM have been limited by its chemical heterogeneity and physical occlusion within the soil matrix (*1, 13*–*19*). Several studies have attempted to fractionate natural soil organic matter into two operationally-defined types - particulate organic matter and MAOM - using traditional density or size separation methods, and to examine their sources, chemical properties, and stability (*20*–*22*). These studies have demonstrated that MAOM exhibits distinct biogeochemical characteristics compared to particulate organic matter, resulting in greater stabilities and longer soil residence time (*10, 20, 23, 24*). However, the inherent complexity of natural soil systems limits further mechanistic investigation into the process that governs their preservation mechanisms in soils.

Because there has been no reproducible method to create a biologically complex, compositionally controlled version of MAOM, many hypotheses about the controls on soil carbon sequestration, such as the roles of minerals and carbon chemistry in MAOM formation and stabilization, have remained experimentally inaccessible. Here, we describe the microbial production and characterization of a novel model material, termed LabMAOM, that makes such mechanistic studies possible. LabMAOM chemistry is both reproducible and tunable on laboratory timescales. We demonstrate LabMAOM can be used to verify existing hypotheses about the nature and behavior of soil carbon and to test previously inaccessible hypotheses on the fundamental controls governing the stability of the soil carbon reservoir.

### LabMAOM reproduces key features of natural MAOM

We chose to microbially synthesize a mimic for natural MAOM because microbes have been implicated in its natural production (7). We sought to biologically replicate the close organomineral associations characteristic of natural MAOM (*14*). We also wanted to ensure that we incorporated similar biomass and mineral components as natural MAOM (*25*–*28*). Our first step towards synthesizing a mimic of natural MAOM – termed LabMAOM – required the identification of a suitable microbial host and a compatible mineralization-facilitating ion (*29*). To enable reproducible and tunable control over biomass composition, we sought a genetically tractable soil microbe capable of producing substantial amounts of biofilm biomass in culture. We selected *Bacillus subtilis* 168, a “generally recognized as safe” organism compatible with agricultural applications, as our chassis (*30, 31*). To increase *B. subtilis* biofilm mass, we deleted the pulcherriminic acid biosynthesis cluster (*32*). We also sought a biocompatible metal ion that could undergo rapid biomineralization while minimally inhibiting biofilm synthesis. We converged on Fe (III), an ion precursor abundant in natural MAOM. To synthesize LabMAOM, we grew *B. subtilis* biofilms in a defined chemical medium (*33*), incubated these biofilms for 10 days at 30 °C with high concentrations of soluble Fe^3+^ as Fe_2_(SO_4_)_3_, and then harvested biomineralized biofilms - called LabMAOM - for downstream characterization and experimentation (Fig. 1a). In ten days, our approach produces 0.6 ± 0.2 g/L of LabMAOM, which is a dark powdery substance.

**Figure 1:**
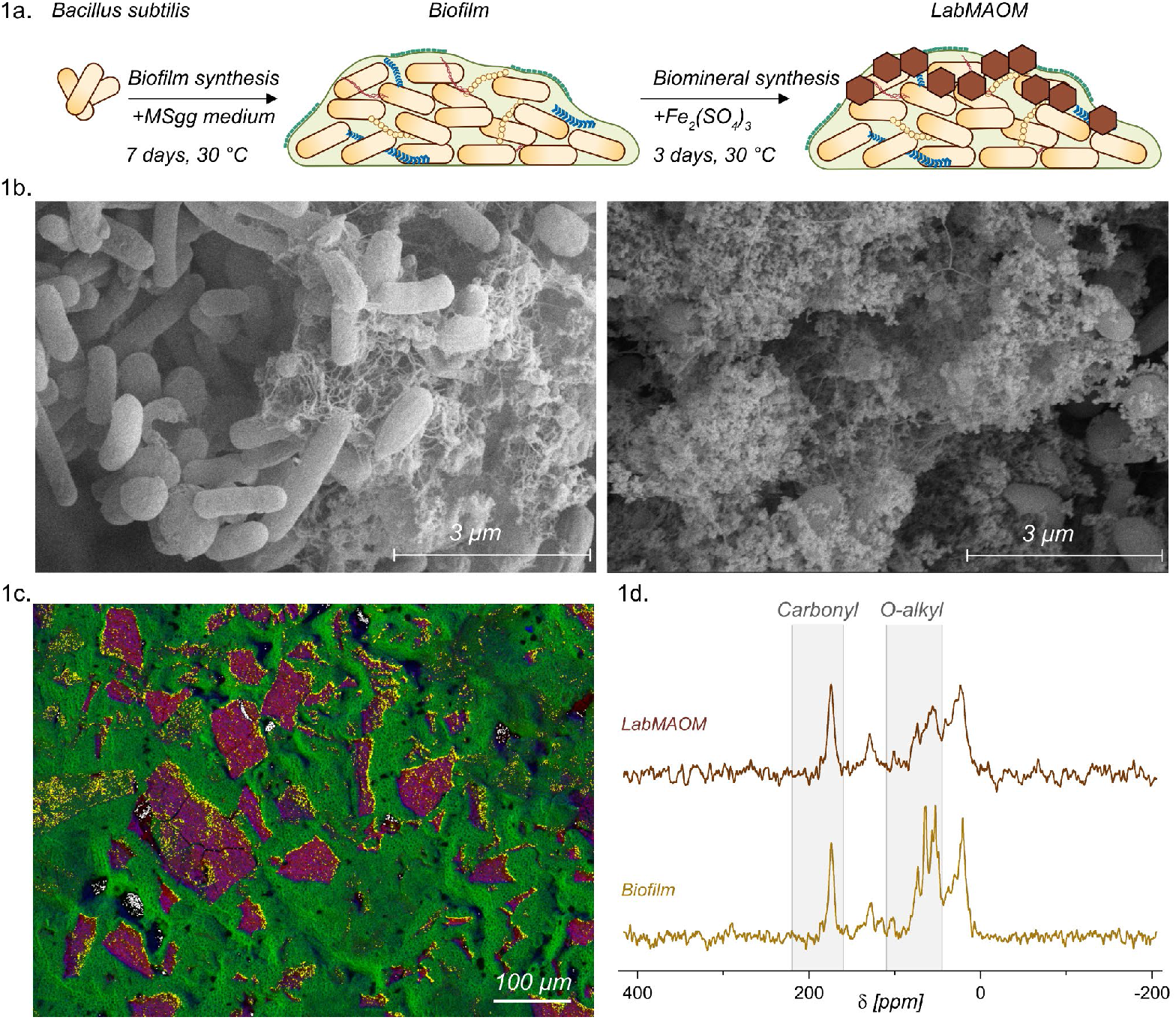
LabMAOM reproduces key features of natural MAOM. **(A)** Schematic representation of LabMAOM synthesis. **(B)** Scanning electron micrographs of unmineralized *Bacillus subtilis* biofilms (left) and LabMAOM (right). Representative LabMAOM micrograph indicates the presence of short-range order iron oxides templated onto biofilm polymers, such as exopolysaccharides. **(C)** WDS of LabMAOM with an Fe (red), C (green), and O (blue) RGB composite overlay. Co-localization of Fe and C is indicated in yellow. Maps indicate the presence of Fe-rich solids intercalated with C and O. **(D)** ^13^C CP/MAS-NMR spectra of unmineralized biofilms and LabMAOM. Spectra exhibit an enrichment of carbonyl and a depletion of O-alkyl moieties in LabMAOM, when compared to a biofilm control. Spectra were normalized to the carbonyl maximum intensity at 173.8 ppm.

Natural MAOM demonstrates key structural and mineralogical features, such as close association of biomass with minerals and presence of short-range order iron oxides (*7, 19, 34*–*37*). To examine the extent to which LabMAOM reproduces these structural and mineralogical features, we probed the morphology and elemental composition of unmineralized *B. subtilis* biofilms and LabMAOM with scanning electron microscopy (SEM) coupled with wavelength-dispersive X-ray spectroscopy (WDS). SEM analysis of the unmineralized *B. subtilis* biofilm (Fig. 1b, left) showed cells connected by a stringy biofilm matrix. These cells and matrices were also present in LabMAOM samples, but they were closely associated with biominerals (Fig. 1b, right). SEM micrographs further reveal that these biominerals, possibly iron oxides, are not well-crystallized and show a lack of long-range order, indicating the presence of short-range order minerals. Furthermore, WDS spectroscopic mapping of LabMAOM (Fig. 1c) showed that the biominerals are indeed composed of iron and oxygen, indicating they are short-range iron oxides. Thus, LabMAOM reproduces key structural and mineralogical features of natural MAOM.

Natural MAOM also has a characteristic chemical signature, an enrichment of carbonyl carbon and depletion of O-alkyl carbon, when compared to surrounding soil organic matter (*38*–*42*). To determine whether LabMAOM reproduces this chemical signature relative to unmineralized *B. subtilis* biofilm controls, we probed both samples using solid-state ^13^C Cross Polarization/Magic Angle Spinning Nuclear Magnetic Resonance (CP/MAS NMR) spectroscopy. Indeed, LabMAOM displays distinct chemical characteristics from unmineralized biofilms (Fig. 1d). In particular, LabMAOM is enriched in carboxyl/carbonyl and depleted in O-alkyl functional moieties, while unmineralized biofilms are enriched in O-alkyl functional moieties. This carbonyl enrichment is a key chemical signature previously observed in natural MAOM that is effectively reproduced in LabMAOM. Taken together, these features show that LabMAOM captures key structural, mineralogical, and chemical features of natural MAOM, yet can be reproducibly synthesized under laboratory conditions.

### Incubation experiments with LabMAOM indicate both mineralogical and environmental controls on carbon preservation

While there is ample empirical evidence highlighting the presence of minerals in persistent soil organic carbon, it remains unclear if mineral association is simply correlated with other parameters, such as occlusion and carbon chemistry, that themselves act mechanistically to control organic carbon persistence (*1, 19, 36, 43, 44*). It has been argued that environmental parameters, such as temperature and ambient labile organic matter, play important roles in controlling soil organic carbon fluxes (*1, 36, 45*–*47*). However, due to the chemical complexity and physical heterogeneity of the soil matrix, it remains challenging to determine the individual contributions of mineral association and environmental parameters to MAOM preservation.

We posited that experiments probing carbon persistence using LabMAOM could improve our understanding of how mineral association and environmental parameters affect natural MAOM persistence. To test the effect of mineral association under conditions of abundant organic matter, we compared the decomposition of unmineralized *B. subtilis* biofilms with that of LabMAOM, both incubated with a model soil degrader, *Pseudomonas putida*, in an organic-rich medium (LB). Decomposition was assessed by monitoring CO_2_ produced in the headspace by *P. putida*. A central challenge in this experiment is distinguishing CO_2_ produced from decomposing *B. subtilis* biomass from CO_2_ derived from the organic carbon present in the *P. putida* incubation medium (LB). We leveraged differences in stable carbon isotopic signatures (δ^13^C) between the medium used to generate *B. subtilis* biomass (MSgg) and the *P. putida* incubation medium (LB) to track decomposition of these distinct carbon pools. We expect that, if *B. subtilis* biomass in LabMAOM is decomposed by *P. putida*, we will observe lower δ^13^C values in the headspace CO_2_, resembling those of the *B. subtilis* biofilm. Conversely, if the biomass in LabMAOM is not degraded, we expect higher δ^13^C values, similar to the headspace CO_2_ from *P. putida*-only controls.

After incubating LabMAOM with *P. putida* in the presence of ambient labile organic matter (LB), we measured the isotopic signature of CO_2_ in the headspace. The δ^13^C of headspace CO_2_ from incubations containing only *P. putida* was significantly higher than from incubations that also contained unmineralized *B. subtilis* biomass (-20.71 ± 0.18 ‰ vs. -22.71 ± 0.53 ‰, Fig. 2b). Furthermore, the δ^13^C of headspace CO_2_ from incubations containing *P. putida* with LabMAOM (-20.55 ± 0.61 ‰) was significantly higher than from incubations that contained *P. putida* with unmineralized *B. subtilis* biomass, and within the same range as incubations that contained only

**Figure 2.**
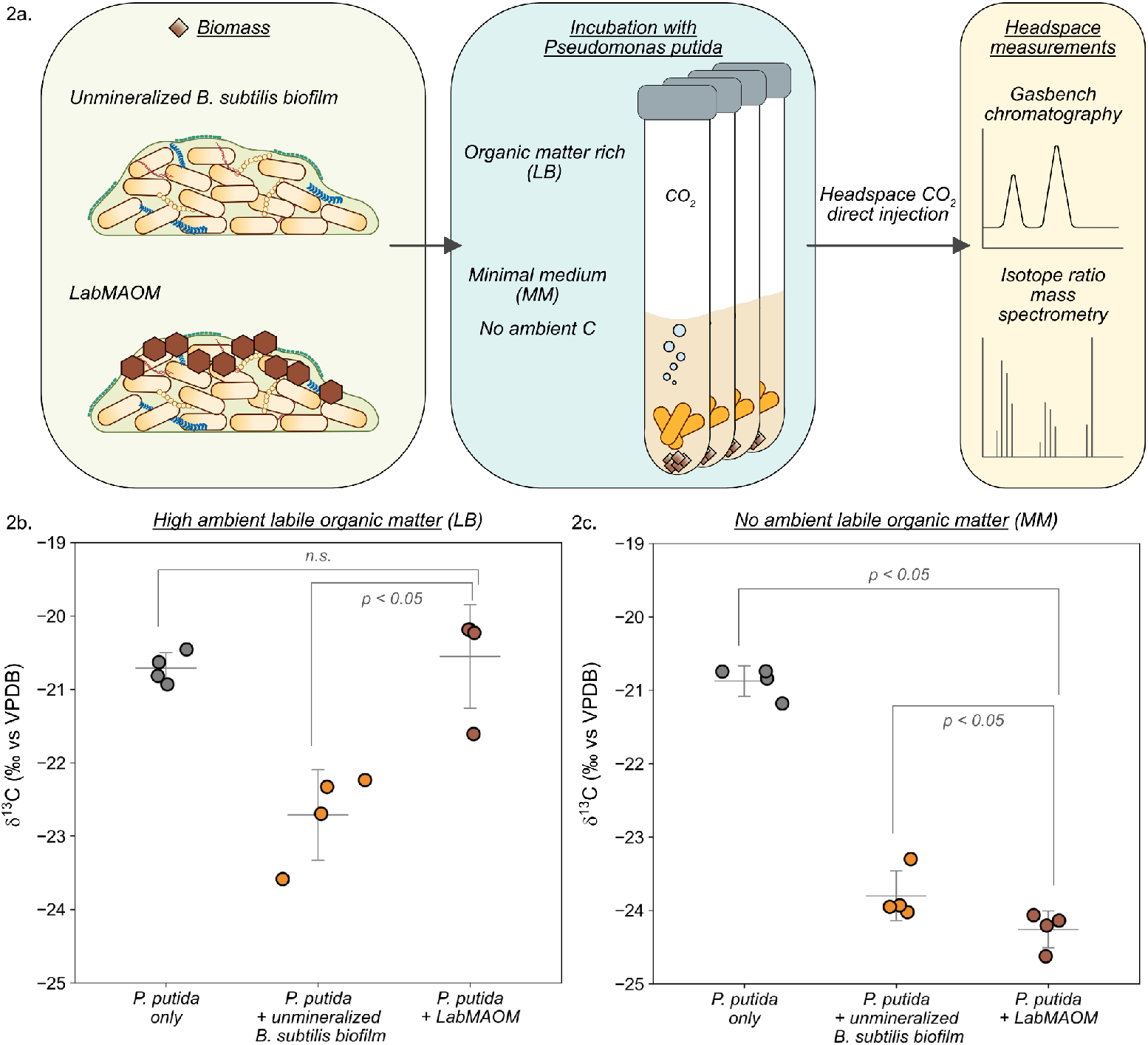
Mineral association and ambient labile organic matter protect LabMAOM from decomposition. **(A)** Schematic of incubation experiments with *Pseudomonas putida* as a degrader. Dried LabMAOM and unmineralized biofilm controls were incubated with *P. putida* in either a rich medium (LB, high ambient labile organic matter) or minimal medium (MM, no ambient labile organic matter) for 7 days at 30 °C. Following incubation, headspace CO_2_ was collected, and δ^13^C was measured using Gasbench-IRMS. **(B)** Headspace CO_2_ δ^13^C measurements from LB incubations of *P. putida* alone (medium only), with unmineralized biofilm, and with LabMAOM. δ^13^C values from LabMAOM incubations were similar to *P. putida*-only controls and significantly higher than unmineralized biofilm, indicating that LabMAOM is protected from degradation by *P. putida* in the presence of minerals and abundant labile organic matter. **(C)** Headspace CO_2_ δ^13^C measurements from MM incubations of *P. putida* alone, with unmineralized biofilm, and with LabMAOM. Here, δ^13^C values from the *P. putida*-only control were significantly higher, while values from both the unmineralized biofilm and LabMAOM incubations were lower, indicating that LabMAOM is degraded by *P. putida* in the absence of ambient labile organic matter, despite the presence of minerals. All samples were measured in biological quadruplicates. Data are presented as arithmetic mean ± standard deviation. Statistical significance was assessed using the non-parametric Mann-Whitney U test.

*P. putida* (Fig. 2b). These measurements showed that δ^13^C values from LabMAOM incubations were similar to those from *P. putida*-only controls and significantly higher than those from incubations containing unmineralized biofilm (Fig. 2b), indicating that the organic carbon in LabMAOM is preserved relative to unmineralized biomass. Together, these results strongly suggest that mineral associations help protect organic carbon under conditions of high ambient organic matter availability.

Next, we sought to test the effect of mineral associations on carbon preservation in the absence of ambient organic matter. To do so, we replaced the *P. putida* incubation medium with minimal medium (MM), which lacks an external carbon source. As before, we measured headspace CO_2_ δ^13^C from the decomposition of *B. subtilis* biomass (MSgg) and *P. putida* in MM to confirm that we could track the decomposition of these distinct carbon pools. Again, we observed that δ^13^C values of CO_2_ from *P. putida*-only incubations were significantly higher than those from incubations containing *B. subtilis* biomass and *P. putida* (-20.87 ± 0.18 ‰ vs. -23.80 ± 0.29 ‰, Fig. 2c), demonstrating that we could resolve which biomass was being decomposed. We then incubated LabMAOM with *P. putida* in MM and measured the isotopic signature of the headspace CO_2_ (Fig. 2c). In this case, headspace δ^13^C values from LabMAOM incubations (- 24.25 ± 0.22 ‰) were similar to those from the unmineralized biofilm control and significantly lower than those from the *P. putida*-only control. These results indicate that in the absence of ambient labile organic matter, minerals do not provide a significant protective effect for the biomass within LabMAOM. More specifically, these findings indicate that LabMAOM remains persistent in environments with high ambient labile organic matter availability, underscoring the role of ambient labile organic matter as a critical environmental control on LabMAOM preservation. Together, these results highlight the joint roles of mineral association and ambient labile organic matter in stabilizing soil organic carbon.

### Carbonyl enrichment is a conserved feature of LabMAOM formation

Extensive empirical evidence demonstrates a tightly-controlled, low C:N ratio for natural MAOM, even in soils with high ambient C content (*48*–*51*). Natural MAOM is also typically enriched in carbonyl C and depleted in O-alkyl C when compared to surrounding soil organic matter (*38, 40, 42*). While these bulk chemical features are widely represented in natural MAOM samples, it remains unclear whether these features are intrinsic to natural MAOM formation. Determining this intrinsic chemical signature is essential to understanding the chemical controls on carbon preserved in organomineral associates.

To synthesize LabMAOM with different bulk chemical compositions, we first grew *B. subtilis* biofilms in MSgg medium with varying glycerol concentrations (0.5%, 1%, 2%, and 5%). This approach yielded a panel of biofilms with modestly distinct yet highly reproducible C:N ratios (Fig. S4) and bulk carbon chemistries (Fig. 3b). Specifically, the C:N elemental ratios varied from 4.941 ± 0.259 at 0.5% glycerol synthesis conditions to 6.346 ± 0.109 at 5% glycerol synthesis conditions (Fig. S4). To test whether natural MAOM formation drives specific chemical features, we used these biofilms as chemically distinct inputs to synthesize LabMAOM and examined their bulk chemical composition using ^13^C CP/MAS NMR spectroscopy. Our analysis revealed that although the chemical composition of input biomass varied substantially, with increasing abundances of O-alkyl moieties (Fig. 3b), the chemical composition of the resulting LabMAOM remained conserved (Fig. 3c, Fig. S5). We also observed characteristic carbonyl enrichment in all LabMAOM samples, regardless of the carbonyl content of their respective input biofilms. Taken together, these results demonstrate that LabMAOM enables testing of hypotheses regarding natural MAOM formation and strongly suggest that carbonyl enrichment and O-alkyl depletion are intrinsic features of natural MAOM formation.

**Figure 3.**
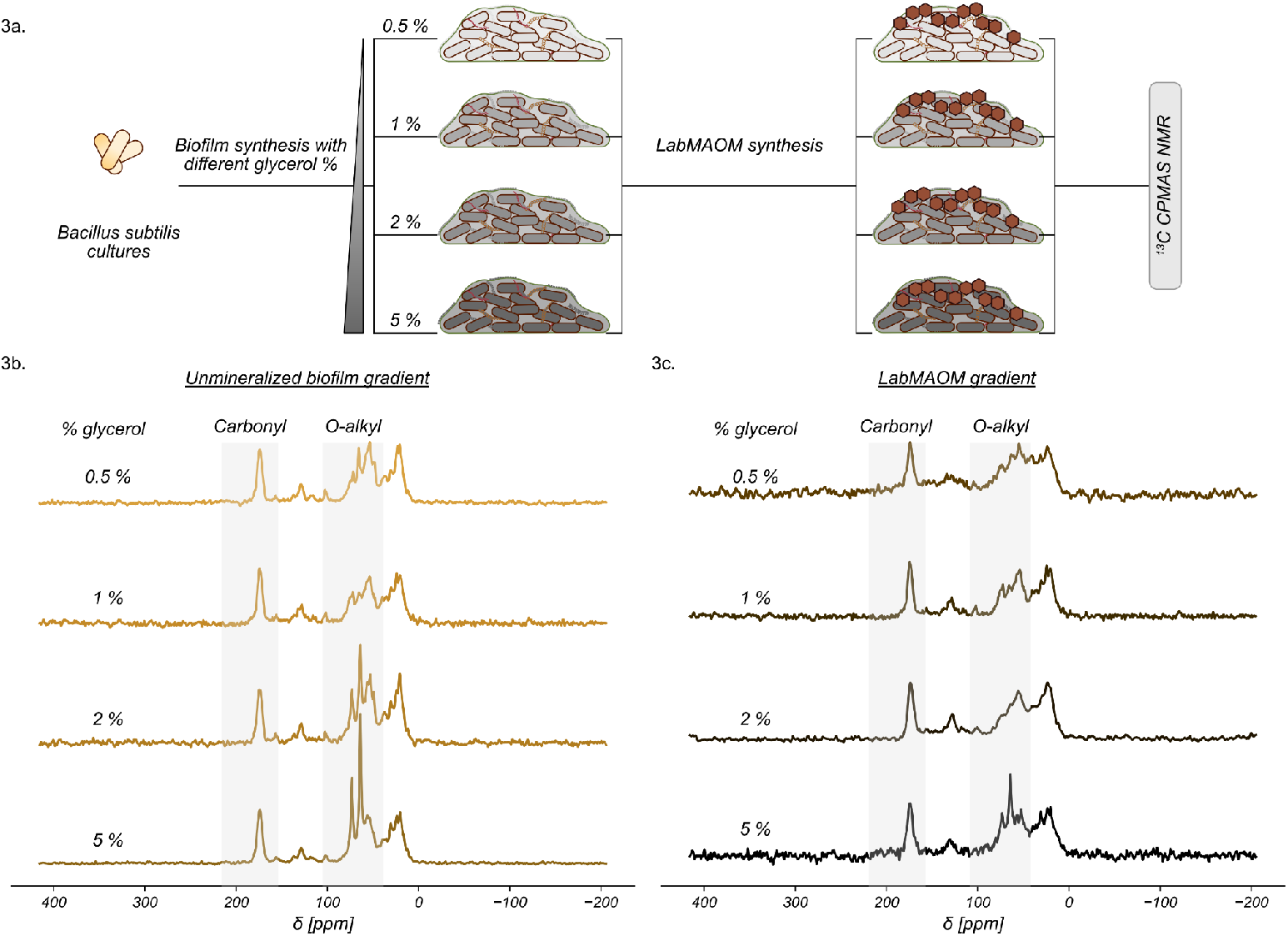
Carbonyl enrichment is a conserved feature of LabMAOM formation. **(A)** Schematic of LabMAOM synthesis under varying carbon conditions. A glycerol gradient was used to generate *B. subtilis* biofilms with differing bulk compositions, which served as templates for LabMAOM synthesis. The grayscale gradient indicates increasing glycerol concentrations during biofilm synthesis, corresponding to higher exopolysaccharide abundance. Bulk chemical composition and functional group distributions were assessed by ^13^C CP/MAS NMR spectroscopy. **(B)** ^13^C CP/MAS NMR spectra of representative unmineralized biofilms synthesized with 0.5%, 1%, 2%, and 5% glycerol. Increasing glycerol concentrations led to a pronounced enrichment in O-alkyl functional groups, indicative of increased exopolysaccharide content. Spectra represent pooled samples from 30 independently synthesized biofilms and are normalized to the carbonyl maximum at 173.8 ppm. **(C)** ^13^C CP/MAS NMR spectra of LabMAOM synthesized using biofilms grown at 0.5%, 1%, 2%, and 5% glycerol as templates. Spectra show depletion of O-alkyl functional groups and a corresponding enrichment of carbonyl moieties, consistent with trends observed in Fig. 1d. Spectra represent pooled samples from 30 independently synthesized LabMAOM preparations and are normalized to the carbonyl maximum at 173.8 ppm.

While this carbonyl enrichment is critical for strong ligand exchange interactions at mineral surfaces (*14*), it is surprising to observe this enrichment in LabMAOM, in which minerals were biomineralized *de novo* rather than from mineral surfaces in soils, as is the case with natural MAOM. However, our results align with previous work on carbon preservation by the coprecipitation of organic matter with short-range order iron (oxyhydr)oxide minerals, indicating that LabMAOM formation is likely a result of coprecipitation (*41, 42, 52*). Additionally, while this carbonyl enrichment has been previously observed in samples of natural MAOM, our work converges on a fundamental chemical composition of natural MAOM that is invariant to input biomass chemistry.

Our work provides a method for the synthesis of a lab standard for natural MAOM, enabling new approaches to key questions on the environmental stabilization of soil organic carbon. We engineered a native soil organism, *Bacillus subtilis* 168, to synthesize LabMAOM and successfully reproduced physical and chemical signatures present in natural MAOM, including close association between microbial biomass and biominerals, the presence of short-range order iron oxides, and the enrichment of carbonyl moieties coupled with the depletion of O-alkyl moieties. We then used LabMAOM in incubation experiments as a model for natural MAOM to test two hypothesized controls governing carbon sequestration in soils: 1) the roles of mineral association and ambient carbon in natural MAOM stabilization, and 2) the role of mineral association in preserving specific carbon functional groups. We determined that mineral association and ambient labile organic matter play synergistic roles in the stabilization of natural MAOM against degradation. More specifically, we demonstrated that, together, mineral association and high ambient labile organic matter protect LabMAOM from degradation, a phenomenon previously observed but not mechanistically explained in samples of natural MAOM.

Additionally, we demonstrated an enrichment of carbonyl and depletion of O-alkyl functional moieties in LabMAOM that is invariant to input biomass chemistry. While this chemical motif has been reported in natural MAOM, our results strongly suggest that this fundamental chemistry is a consequence of natural MAOM formation. While it has been previously observed that natural MAOM maintains a tightly-regulated C:N ratio (*43, 48–51*), less is known about the processes that control the ancillary chemical speciation of carbon in natural samples. Our work hints at a mineralogical control on the chemical composition of natural MAOM, which we speculate may be a critical control on the types of organic carbon that remain stabilized in soils for millennia. Further experiments with LabMAOM could shed light on whether this conserved chemical composition is representative of samples synthesized from various litter sources, such as leaves and agricultural waste.

This work addresses a fundamental uncertainty in our understanding of carbon stabilization in the Earth system and opens new doors for carbon sequestration in a changing climate. A central uncertainty in our understanding of soil organic carbon stabilization is the relative importance of mineral association versus environmental controls on organic carbon degradability. Without a standard material, it has not been previously possible to perform reductionist experiments that isolated these single parameters in a controlled manner. LabMAOM will enable a systematic exploration of how other environmental variables, such as soil pH, temperature, and mineralogy, influence soil carbon sequestration (*1, 22, 36, 46, 53, 54*). LabMAOM will also elucidate chemical and mineralogical controls on soil carbon stabilization, such as the relative roles of the earth-abundant metals Fe, Al, Ca, and the presence of ambient N and P. This information will be immediately economically relevant in areas as diverse as carbon removal through enhanced rock weathering, to improvements in crop amendments to increase soil performance while expanding our fundamental understanding of controls on the global carbon cycle. Finally, as the synthesis strategy for LabMAOM can be transferred to an autotrophic chassis organism, our work provides a path towards the synthesis of a carbon-negative soil amendment.

## Acknowledgments

The authors acknowledge instrumental support from Dr. Gelu Costin.

## Funding

Moore Foundation Grant # 7524

Welch Foundation Award # C-2130-20230405

Office of Naval Research (ONR) Grant # N00014-21-1-2362

U.S. Department of Energy Office of Science Energy Earthshot Initiative, as part of the LLNL

Terraforming Soils EERC under Award # SCW1841.

## Author contributions

Conceptualization: SS, CAM, CMAF

Data Curation: SS, XG, TS

Formal Analysis: SS, XG, TS

Investigation: SS, XG, TS

Methodology: SS, XG, TS

Software: SS

Visualization: SS

Funding Acquisition: CAM, CMAF

Project Administration: CAM, CMAF

Resources, Supervision & Validation: CAM, CMAF

Writing, original draft: SS, CAM, CMA-F

Writing, review & editing: SS, XG, TS, CAM, CMAF

## Data and materials availability

The Bacillus subtilis Δ*cypX-yvmC* strain is available upon request with completion of an MTA. All data are available in the main text or the supplementary materials.

Code repository: https://github.com/ssridhar19/LabMAOM

## Supplementary Materials

Materials and Methods

Supplementary Text

Figs. S1 to S5

Tables S1 to S4

