## Supplementary Materials for "Mechanisms of soil carbon preservation illuminated by model mineral-associated organic matter"

**Supplementary Material** for “Mechanisms driving soil carbon preservation illuminated by model mineral-associated organic matter”

### Materials and Methods

Media and growth conditions

All strains used in this study were constructed in *Bacillus subtilis* 168 and are described in **Table S1**. *B. subtilis* 168 was purchased from ATCC (str. *Bacillus subtilis* (Ehrenberg) Cohn, #23857). Routine culture of *B. subtilis* 168 was performed in LB (Sigma-Aldrich, L3522). Biofilms were synthesized in minimal medium MSgg: 5 mM $KPO_4$ buffer (pH 7, for 1 M $KPO_4$ in 100 mL: 9.343 g $K_2HPO_4$ and 6.309 g $KH_2PO_4$), 100 mM MOPS (pH 7), 2 mM $MgCl_2$, 700 uM $CaCl_2$, 50 uM $MnCl_2$, 50 uM $FeCl_3$, 1 uM $ZnCl_2$, 2 uM thiamine, 50 ug/mL tryptophan, 50 ug/mL phenylalanine, 0.5% glutamate, 0.5% glycerol (*1*).

All LabMAOM incubation experiments were performed with *Pseudomonas putida* str. F1 as a model soil degrader. Freshly-picked colonies of *P. putida* were inoculated in 1 mL LB and incubated at 30 °C, 250 rpm for 12 h. This pregrown culture was washed three times with autoclaved, distilled $H_2O$ before diluting 1:100 in minimal medium (MM) for incubation experiments.

MM (pH 7, per L): 1 mL 20% $MgCl_2.6H_2O$, 0.1 mL SL7 (per 100 mL: 1 mL 25% HCl, 0.07 g $ZnCl_2$, 0.1 g $MnCl_2.4H_2O$, 0.06 g $H_3BO_3$, 0.2 g $CoCl_2.6H_2O$, 0.02 g $CuCl_2.2H_2O$, 0.02 g $NiCl_2.6H_2O$, 0.04 g $NaMoO_4.2H_2O$), 4.68 g NaCl, 1.49 g KCl, 1.07 g $NH_4Cl$, 0.43 g $Na_2SO_4$, 0.03 g $CaCl_2.2H_2O$, 0.23 g $Na_2HPO_4.12H_2O$, 0.005 g Fe(III)$NH_4$ citrate. Filter sterilize through a 0.22 um filter.

DNA manipulation

All plasmids used for genome editing (**Table S2**) were constructed using Gibson assembly. Fragments for Gibson assembly (**Table S3**) were amplified using PhantaFlash polymerase (Vazyme, P520) according to manufacturer recommendations, with reaction-specific PCR conditions summarized in **Table S4**. PCR products were purified from agarose gels using the GenCatch Gel Extraction Kit (Epoch Life Science, 2260050) and assembled using the NEBuilder HiFi assembly system (New England Biolabs, E2621). Assembled plasmids were transformed and maintained in NEB 10-beta competent *E. coli* (New England Biolabs, C3019) and purified from *E. coli* using the QIAprep Spin Miniprep Kit (Qiagen, 27104). All plasmids were sequenced using long-read Nanopore sequencing (Plasmidsaurus).

Competent cell preparation and transformation

Competent cell preparation and transformation was performed as described in Jayaraman et al with minor modifications (*2*). *B. subtilis* 168 was streaked onto LB plates and grown overnight at 37 °C. A single colony was inoculated in 1 mL of competence medium (0.5% glucose, 1.4% $K_2HPO_4$, 0.6% $KH_2PO_4$, 30 nM Na-citrate tribasic dehydrate, 0.2% casein hydrolysate, 84 mM $NH_4$Fe(III) citrate, 3 uM $MgSO_4$) and incubated at 37 °C, 250 rpm for 3.5 h. 300 ng of plasmid

DNA was added to 300 uL of freshly prepared competent cells and incubated at 30 °C, 250 rpm for 4 h. Entire transformant volume was plated onto LB with antibiotics and incubated at 30 °C for 12 h or till colonies were visible.

Pulcherriminic acid deletion

To overcome the inherent limitations on biofilm mass synthesis, we deleted the pulcherriminic acid biosynthesis cluster. Previous work has shown that deleting pulcherriminic acid biosynthesis results in evergrowing *B. subtilis* biofilms (*3*). We hypothesized that this feature of *B. subtilis* biofilms can be leveraged to synthesize more biofilm mass, which would subsequently result in more LabMAOM.

To test our hypothesis, we deleted the *cypX-yvmC* cluster from *B. subtilis* using a two-step double-homologous recombination protocol. In the first step, we transformed naturally-competent *B. subtilis* 168 cells with plasmid pSS172 (**Table S1**) to facilitate a scarred knockout of *cypX-yvmC*, with *mScarlet-I* for screening and *ermB* for selection on 1 ug/L Erythromycin. Bright pink colonies were inoculated for competent cell preparation as described previously. Naturally competent Δ*cypX-yvmC*::*mScarlet-I-ermB* cells were transformed with plasmid pSS167 (**Table S1**) to delete the *mScarlet-I-ermB* scar and produce a scarless genomic knockout. As pSS167 has *sfGFP* for screening and a temperature-sensitive oriV for plasmid curing, we selected colonies that displayed bright red and green fluorescence, and cured pSS167 on LB plates at 37 °C for 12 h. Colonies were screened for the absence of *cypX-yvmC* by colony PCR with oligos oSS1259 and oSS1260.

Bacterial and biofilm growth conditions

To grow *B. subtilis* biofilms, cells from frozen glycerol stocks were first streaked onto LB plates and incubated at 30 °C for 12 h. Single colonies were inoculated into 3 mL of LB and cultured at 30 °C, 250 rpm for 8 h. To retain peptides for quorum sensing induction, unwashed cells were inoculated directly into biofilm synthesis medium (MSgg) at a 10x dilution. Biofilms were grown without shaking at 30 °C in covered 12-well plates (Corning, #3513) with 3 mL MSgg in each well. After incubating for 7 days, 300 uL of 12.5 mM $Fe_2(SO_4)_3$ (399.88 g/mol, Sigma Aldrich) stock solution in $ddH_2O$ was added to each well for a final $Fe_2(SO_4)_3$ concentration of 1.25 mM. Samples were covered in foil and incubated at 30 °C for 3 more days to promote biomineralization.

Samples were harvested, centrifuged at 4000 rpm, 22 °C, 15 min to pellet biofilms, and lyophilized at standard vacuum conditions (0.013 mbar, -80 °C) for 24 h. Prior to lyophilization, a fourth of each sample was reserved for scanning electron microscopy-wavelength dispersive spectrometry (SEM-WDS).

Scanning electron microscopy and Wavelength Dispersive Spectrometry

Harvested samples were preserved in 1 mL ice cold Karnovsky's fixative and stored in 4 °C for 24 h. The fixative was washed out thrice with 0.1 M HEPES (pH 7.2) for 10 min each. Washed samples were dehydrated in a gradient alcohol series, with two washes at each step of the concentration gradient (25%, 50%, 70%, 95%, 100% EtOH). Samples were further incubated in 100% EtOH for 1 h and dried using a Critical Point Dryer to maintain bacterial cell structures. Dehydrated and dried samples were mounted onto a 12.7 mm aluminium stub and coated with 10 nm carbon. Samples were imaged in a ThermoFisher Apreo electron microscope at 5 kV with the T1 detector.

Wavelength Dispersive Spectrometry (WDS) element mapping was acquired using 15 kV accelerating voltage, 10 nA beam current, and 50-200 ms dwell time, using stage mode scanning. Deadtime correction was applied for every mapped element. Each element map was subsequently quantified employing the standards: graphite for C, olivine for Si, Mg and Fe, chromite for Cr, plagioclase for Na and Ca, pyrite for S, fluorapatite for F, tugtupite for Cl. Data was extracted using SPView software and processed using a custom Python script (edx-modified.py).

Elemental analysis

Elemental combustion analysis was performed on a Costech 4010 CHNS/O Elemental Analysis System to determine C and N composition. Samples were weighed in Sn capsules (5 x 9 mm, Costech) with weights averaging ~ 0.3 mg. Acetanilide was used as a standard (0.05 mg - 3.0 mg), with phenylalanine (~ 1.0 mg) as a quality control internal standard. All samples were run in three biological replicates. Data analysis was conducted within the EA software with a linear fit calibration for nitrogen and a quadratic fit for carbon. Standard curves were generated using a custom Python script (EA-calibration.py).

Bulk carbon isotope analysis

The $\delta^{13}C$ measurements of carbon sources for incubation experiments were performed on an Elemental Analyzer (EA, Flash HT Plus)-Isotope Ratio Mass Spectrometer (IRMS, Thermo Delta V) system. Sample preparation is similar to the aforementioned Elemental Analysis. Produced $CO_2$ gas through combustion (at 1020°C) within EA was transferred to the IRMS for isotope analysis. IAEA CH3 was used as a working standard for calibration purposes. All samples were duplicated or replicated. Analytical precision is 0.2 ‰ or better. Isotopic elemental analysis plots were generated using a custom Python script (isotope-baseline.py, **Fig. S2**).

Incubation experiments and headspace $CO_2$ isotopic analysis

Twelve mg, carbon-normalized amount of biofilm controls and LabMAOM were incubated with P. putida cultures at 30°C for 7 days in gastight borosilicate glass vials (Labco, E2860-100). After 7 days, the δ13C of the headspace $CO_2$ gas was measured using a Gasbench (GasBench II, ThermoFisher)-Isotope Ratio Mass Spectrometer (IRMS, Thermo Delta V). For each sample, the headspace gas (air + respired $CO_2$) was flushed with streams of helium gas at 30°C into the Gasbench for purification. Final $CO_2$ was then delivered to the IRMS for isotope analysis. Quantitative calibration was achieved by acidifying known masses of NaHCO3 (ThermoFisher) and measuring released headspace $CO_2$. Isotopic calibration was achieved by running acidified USGS-44 and IAEA-603 carbonate isotopic standards together with each batch of the samples. No acid was added to experimental vials to avoid any changes in chemistry in subsequent experiments. Apparent isotope difference between $CO_2$ dissolved in solution (2-3 cc) and headspace $CO_2$ is assumed to be negligible. Data plots were generated using a custom Python script (gasbench.py).

$^{13}$C Solid-state NMR

Bulk chemical composition was determined using solid-state $^{13}$C CP/MAS NMR (*4, 5*). Samples were analyzed on a Bruker Avance 200 MHz NMR spectrometer (50 MHz $^{13}$C resonance frequency) equipped with a 4 mm MAS probe and spun at 12 kHz. A modified multiCP/MAS pulse sequence was used to improve quantitative assessment of organic carbon pools (*6*). Chemical shifts were calibrated using a glycine spin-counting standard (*7*).

Spectra were integrated across key chemical shift regions: 0–45 ppm (alkyl-C), 45–110 ppm (O/N-alkyl-C), 110–160 ppm (aryl-C), and 160–220 ppm (carboxyl-C) (*8*). To estimate the relative abundance (%) of biochemical classes (lipids, lignins, proteins, carbohydrates), spectra and C:N ratios were processed using a molecular mixing model (MMM) (*4*). Fits with model errors > 5% were excluded. Spectra were generated using a custom Python script (NMR-plots.py, NMR-spectraldecomp.py).

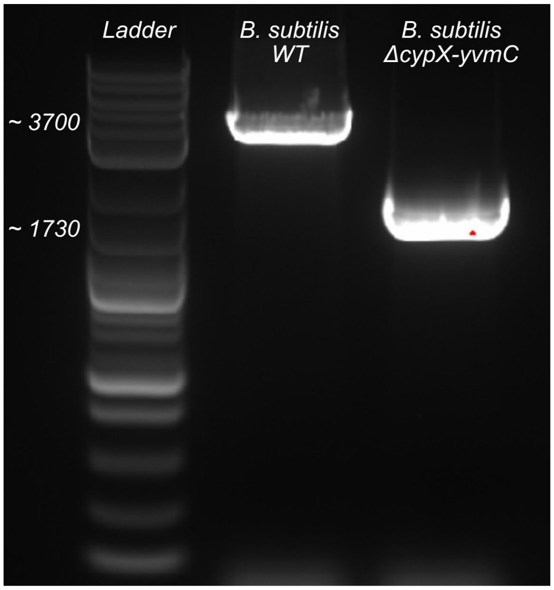


**Fig. S1.**

Image of agarose gel electrophoresis of colony PCR results demonstrating a clean knockout of the *cypX-yvmC* gene cluster. From left to right: DNA ladder (NEB 1 kb Plus, #N0550S), *B. subtilis* wildtype (WT), and *B. subtilis* Δ*cypX-yvmC*.

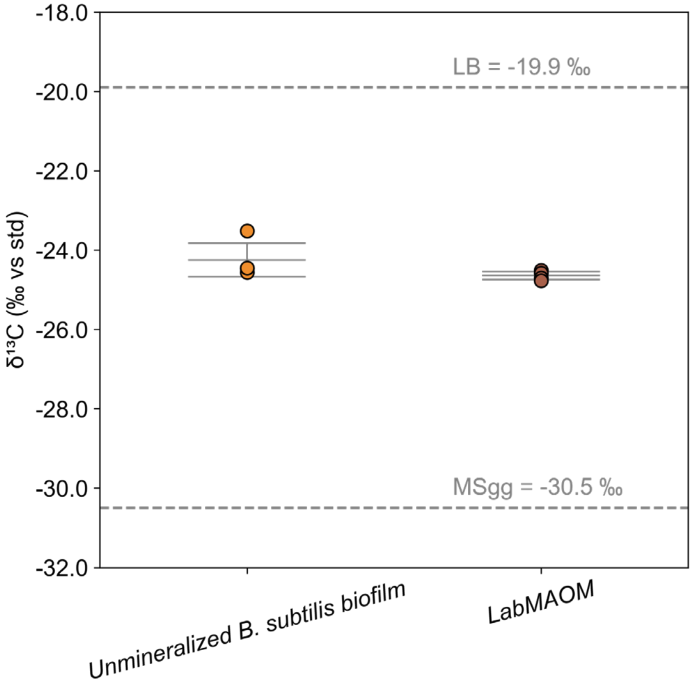


**Fig. S2.**
$^{13}C$ Isotopic elemental analysis for unmineralized *B. subtilis* biofilms and LabMAOM. Gray dotted lines indicate LB (-19.9 ‰) and MSgg (-30.5 ‰) baseline $^{13}C$ fractionation. Summary statistics depict the arithmetic mean and standard deviation.

<insert page break then Fig. S3 here>

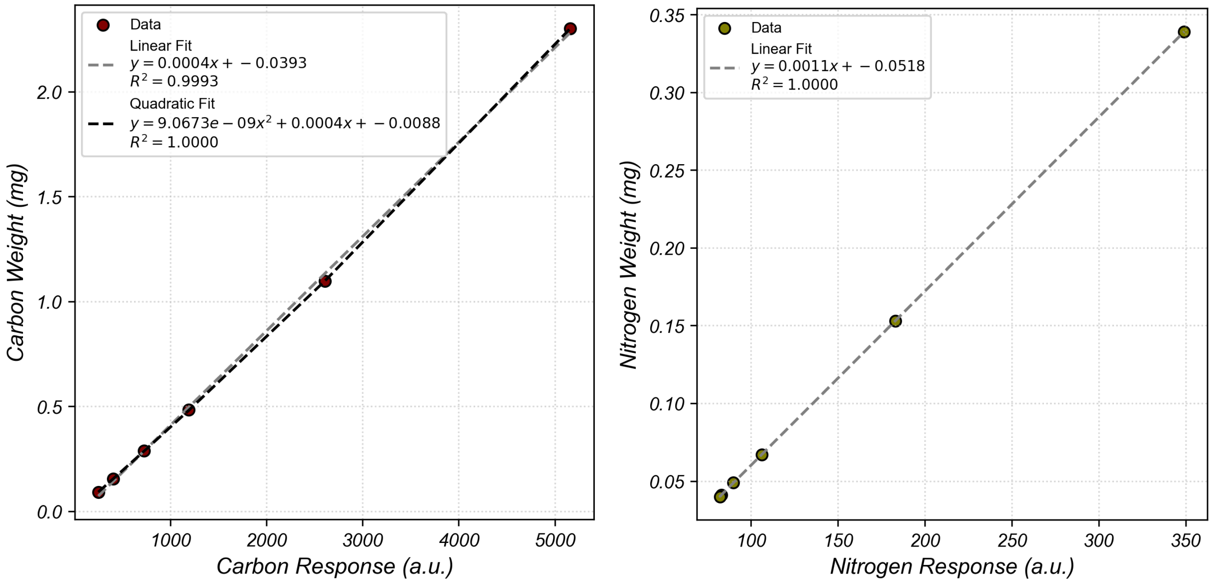


**Fig. S3.**

EA calibration curves for Carbon (left) and Nitrogen (right) generated from a 6-point Acetanilide standard. Carbon data was fitted to both linear and quadratic equations, of which quadratic was selected for data analysis. Nitrogen data was fitted to a linear equation. Fit equations were generated using a custom Python script (EA-calibration.py).

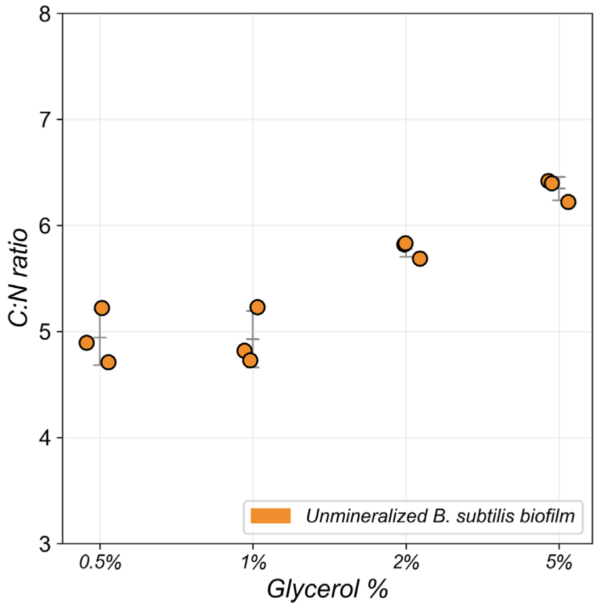


**Fig. S4.**
Bulk C:N ratio of unmineralized biofilms synthesized on a glycerol gradient (0.5% – 5%). Data demonstrate a highly conserved C:N ratio (4.941 ± 0.259 at 0.5% glycerol synthesis conditions to 6.346 ± 0.109 at 5% glycerol synthesis conditions), with a modest increase in C:N ratio with increase in initial glycerol concentration.

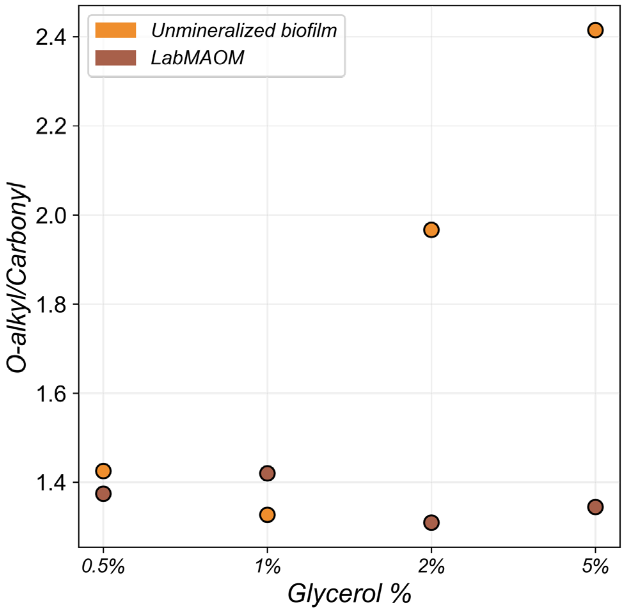


**Fig. S5.**
O-alkyl/Carbonyl ratios of unmineralized biofilms and their corresponding LabMAOM synthesized on a glycerol gradient (0.5% – 5%). O-alkyl/Carbonyl ratios were obtained by normalizing peak areas resulting from spectral decomposition of spectra in Fig. 3b (for unmineralized biofilms) and Fig. 3c (for LabMAOM). Data demonstrate an increase in O-alkyl/Carbonyl ratios with increasing glycerol concentrations for unmineralized biofilms, but a corresponding increase was not observed for LabMAOM samples.

**Table S1.**

Strains used in this study.

| Strain | Purpose |
|---|---|
| *B. subtilis* 168 | Background strain |
| *B. subtilis* 168 Δ*cypX-yvmC* | *B. subtilis* 168 with the pulcherrimin biosynthetic cluster knockout |
| *Pseudomonas putida* F1 | Model degrader organism for incubation experiments |

**Table S2.**

Plasmids used in this study.

| Plasmid | Purpose |
|---|---|
| pSS172 | First step, scarred knockout of *cypX-yvmC*, with *mScarlet-I* for screening and *ermB* for selection on Erythromycin |
| pSS167 | Second step, scarless knockout of *mScarlet-I* and *ermB* |

**Table S3.**

Oligonucleotides used in this study.

| Primer ID | Sequence | Purpose |
|---|---|---|
| oSS1283 | agaaactgcataaaaaaaaggaaagggcctcgtgatacgc | Fragment amplification for Gibson assembly of pSS172 |
| oSS1284 | ctatttcaaacgcctcattattcataaatcagacaaaacttttctcttg | |
| oSS1285 | agttttgtctgatttatgaataatgaggcgtttgaaatag | |
| oSS1286 | gcgtatcacgaggccctttccttttttttatgcagtttctc | |
| oSS1251 | agaaactgcataaaaaaaaagctaacggggcaggttagtga | Fragment amplification for Gibson assembly of pSS167 |
| oSS1252 | ccagaatttctgtctaaaagcttaaaccagttttcgctgg | |
| oSS1253 | ccagcgaaaactggtttaagcttttagacagaaattctggacgtaaa | |
| oSS1254 | ctatttcaaacgcctcattattaagcatgcgtattgggcg | |
| oSS1255 | cgcccaatacgcatgcttaataatgaggcgtttgaaatag | |
| oSS1256 | aagttttaggggtgaatgagtagaattccaaaggtctctc | |
| oSS1257 | gagagacctttggaattctactcattcacccctaaaactt | |
| oSS1258 | tcactaacctgccccgttagcttttttttatgcagtttct | |
| oSS1259 | gctttcacatcaatttgagg | Knockout colony PCR verification |
| oSS1260 | agtgttgacgtcaactgctc | |

**Table S4.**

PCR conditions used in this study.

| Fragment ID | Primer ID | Template | Ta (°C) | Product size (bp) |
|---|---|---|---|---|
| 172-1 | oSS1283, oSS1284 | Non-replicative oriV backbone | 61.8 | 3517 |
| 172-2 | oSS1285, oSS1286 | *B. subtilis genomic DNA* | 50 | 3672 |
| 167-1 | oSS1251, oSS1252 | Replicative, temperature-sensitive oriV (rep E194) backbone | 55.6 | 3495 |
| 167-2 | oSS1253, oSS1254 | | 60.5 | 2305 |
| 167-3 | oSS1255, oSS1256 | *B. subtilis genomic DNA* | 48.4 | 840 |
| 167-4 | oSS1257, oSS1258 | *B. subtilis genomic DNA* | 47.7 | 840 |
| Colony PCR | oSS1259, oSS1260 | *B. subtilis colonies* | 52.7 | 3711 (wildtype)<br>1731 (knockout) |
